# β-adrenergic receptors modulate CA1 population coding during cumulative spatial memory formation and updating

**DOI:** 10.1101/2025.07.07.663515

**Authors:** Ninad Shendye, Josué Haubrich, Jens P Weber, Denise Manahan-Vaughan

**Affiliations:** Ruhr-University Bochum, Medical Faculty, Department of Neurophysiology

**Keywords:** mouse, *in vivo* calcium imaging, learning and memory, hippocampus, neuronal ensemble, adrenergic

## Abstract

Hippocampal neuronal ensembles are likely to support the acquisition, stabilization and updating of spatial experience. Spatial learning is typically cumulative, but little is known about how neuronal ensembles are manifested during this process. Here, we used wide-field Ca^2+^-imaging to monitor CA1 pyramidal cells during cumulative item-place learning in adult male CBA/CaOlaHsd mice. In control mice, initial learning prompted activity in a population of CA1 neurons, some of which re-appeared during re-exposure to the same item-place configuration 60 min after 1^st^ exposure. Item-place reconfiguration (60 min later) caused a change in population dynamics as reflected by alterations in neuronal recruitment and reactivation patterns. Place cell-like properties, population burst activity, and functional connectivity were consistent with the encoding and updating of item-place memory.

To examine the role of noradrenergic neuromodulation on these processes, we pharmacologically antagonized β-adrenergic receptors(β-AR) prior to the 1^st^ item-place exposure. This led to reduced cellular recruitment, disrupted ensemble reactivation, reduced spatial tuning, dampened population bursts, and altered functional connectivity within neurons. This was accompanied by impaired spatial learning compared to controls. Our results reveal the population activity of CA1 neurons during item-place learning and show that β-AR support memory function by influencing both neuronal and network-level dynamics.

## INTRODUCTION

Memory formation and updating rely on the hippocampus, particularly the CA1 region, which integrates spatial and contextual information to support flexible memory representations^1^. The interplay between cellular activity and population-level synchrony within CA1 is essential for encoding new experiences and incorporating them into existing memory frameworks^2–4^. These processes are tightly regulated by neuromodulators, with the noradrenergic (NA) system playing a central role in modulating synaptic plasticity and network dynamics via β-adrenergic receptors^5–8^. However, the mechanisms by which β-adrenergic signaling orchestrates these processes within hippocampal circuits during learning remain poorly understood.

Neuronal ensembles activate in a coordinated manner during memory encoding^9^, and their reactivation supports memory recall^4,10,11^. The hippocampus is central to these processes, mediating memory encoding, consolidation, recall, and updating^12–14^. Within the hippocampus, CA1 neurons are spatial contexts to create cognitive representations of space^2^. During memory encoding and consolidation, CA1 neurons exhibit periodic bursts of synchronous activity^3,15,16^, and the reactivation of neurons that participated in these bursts during encoding has been linked to memory updating^17^. Although electrophysiological recordings from place cells^18^ as well as loss and gain-of-function studies of CA1 pyramidal cells^19^ have demonstrated the necessity of population activity for the mapping of space, as well as for the acquisition, retrieval and updating of spatial memories, little is known about how population dynamics in terms of ensemble activity and neuronal connectivity are engaged during cumulative spatial learning. Multi-unit electrophysiological recordings from neurons during associative learning have revealed transregional co-activity of neurons in cortex and subcortex^20^, whereas recordings from place cells have revealed how context and motivation influences place fields^18,21,22^. However, not all CA1 pyramidal cells are place cells^23^ and neuronal population dynamics can reflect a multitude of other forms of information processing such as stimulus-dependent sensory integration^24^, behavioral/affective state^25^ and temporal experience^26^. Observing the activity of pyramidal cells during cumulative learning and/or information updating can thus, provide novel insights into functional segregation within the CA1 region and how neuronal ensembles are modified by experience^27^. Novel approaches, such as wide-field Ca^2^-imaging of hundreds of CA1 neurons offer an approach through which neuronal ensembles can be monitored in real-time in behaving rodents^28^.

Synaptic plasticity enables the encoding of information by altering the strength of synaptic transmission between neurons^29^. β-adrenergic receptors are densely expressed in the hippocampus^5^ and play a central role in promoting synaptic plasticity in the CA1 region^30–34^. The importance of β-adrenergic receptors for spatial learning and memory has been demonstrated across diverse rodent tasks. Moreover, noradrenergic neuromodulation can alter the location preference of place cells^35^ and neuronal excitation in the hippocampus^36^. Yet, the influence of β-adrenergic receptors on CA1 neuronal ensembles during memory encoding and recall remains unclear.

In this study, we leveraged wide-field Ca^2+^-imaging to investigate how neuronal ensembles develop, stabilize and/or change during item-place learning and the updating of item-place information in mice. In addition, we examined the extent to which β-adrenergic receptor antagonism modulates hippocampal CA1 neuronal activity during these learning events. Our findings reveal the engagement of CA1 neurons, during information acquisition and updating item-place information, at multiple levels including neuronal recruitment, population activity, functional connectivity and place-field mapping. These patterns reflected information acquisition, stabilization and updating. Moreover, we show that β-adrenergic antagonism impaired item-place memory and that this disruption is accompanied by changes in neuronal activity and neuronal ensembles.

## RESULTS

### Pharmacological inhibition of β-adrenergic receptors impairs item-place learning

For this study, we used a cumulative item-place learning and information updating task that is known to induce hippocampal synaptic plasticity and is regulated by β-adrenergic receptors^31^. We monitored both behavioral performance and neuronal Ca^2+^ activity when mice were first exposed to a novel item-place configuration, were re-exposed to the same configuration 1 h later, and then experienced a changed of item-place configuration after a further 60 min. To examine the role of β-adrenergic receptors in these learning events, we administered the antagonist propranolol systemically before task initiation.

One day before novel item-place exposure, animals were habituated to the arena and baseline recordings of Ca^2+^ activity were obtained. On the test day, mice received an intraperitoneal (i.p.) injection of either vehicle (0.1 ml/g body weight), or the β-adrenergic receptor antagonist, propranolol (20 μg per gram of bodyweight at a volume of 0.1 ml/g). Thirty minutes later, the item-place task commenced, consisting of three sessions separated by 60-minute intervals comprising: **Session 1**: Novel item-place Exposure (10 min), in which animals explored two distinct novel objects that were inserted into the recording chamber; **Session 2**: item-place re-exposure (10 min), whereby animals were exposed to the same object configuration; and **Session 3**: item-place reconfiguration (10 min), in which one object’s position was changed relative where it had been in the 1^st^ and 2^nd^ sessions (**Figure 1A**).

**Figure 1:**
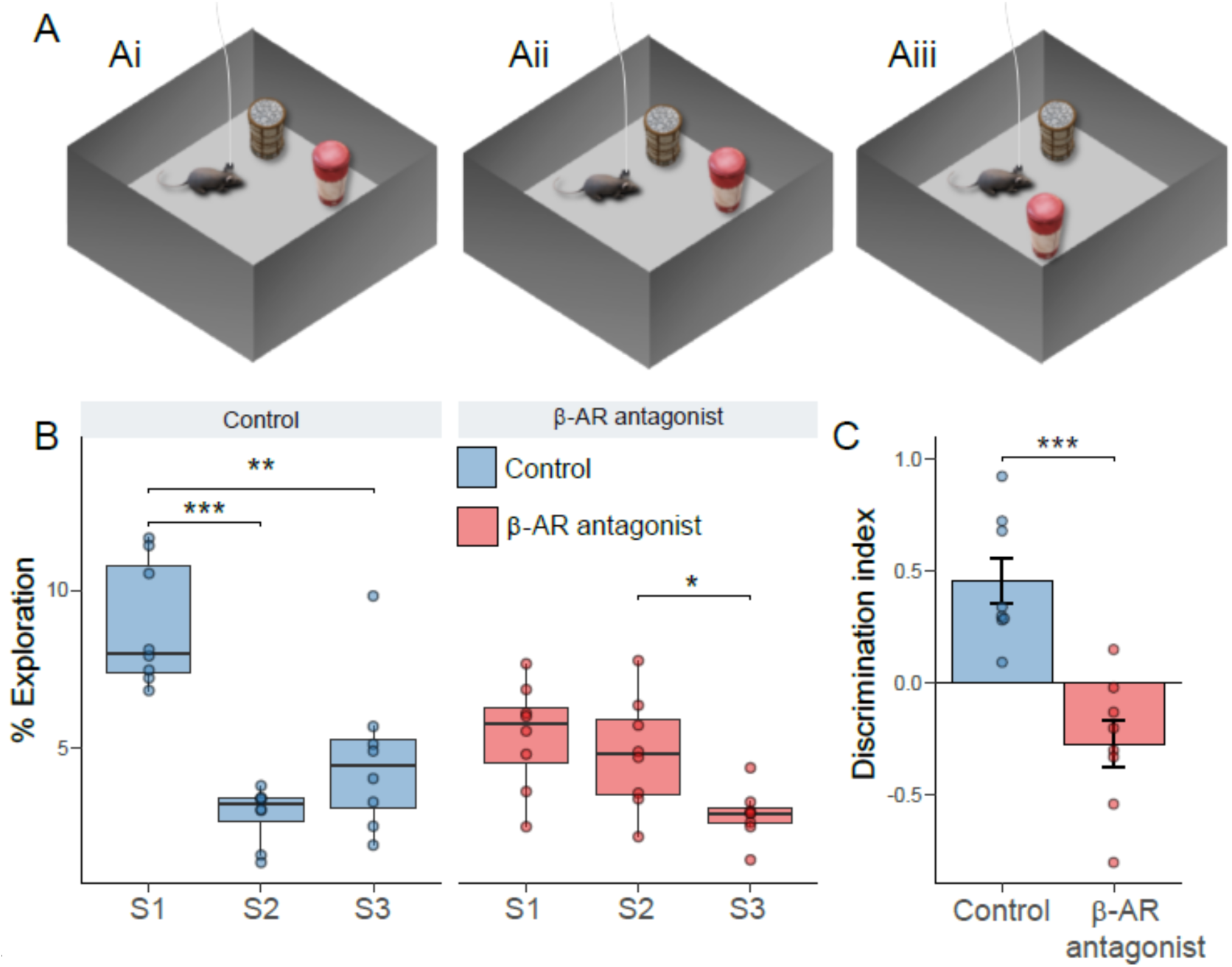
β-Adrenergic receptor antagonism impairs the encoding of item-place associations. **A)** Schematic representation of the experimental design for the item-place learning task consisting of (Ai) session 1 (**S1**, novel item-place configuration); (Aii) session 2 (**S2**, re-exposure to the same item-place configuration) and (Aiii), session 3 (**S3**, displacement of one of the now familiar objects to a different location) **B)** Object exploration times across experimental phases S1 (novel exposure), S2 (re-exposure), S3 (novel configuration). **C)** Discrimination index indicating exploration of displaced versus stationary objects during S3. Boxplots show the median and interquartile range (B), and bars (C) represent the mean ± SEM. Individual data points are overlaid to illustrate the distribution within each group. Asterisks indicate statistical significance at \**p* < 0.05, \*\**p* < 0.01, \*\*\**p* < 0.001. N = 8 per group.

In vehicle-treated animals, the time spent exploring objects varied significantly across sessions (Repeated-measures ANOVA; F2,23 = 24.21, *p* < 0.001), with a decrease evident during session 2 and session 3, compared to session 1 (*p* < 0.05) (**Figure 1B**). During session 3, discrimination indices ^37^ were calculated to determine how well mice distinguished between the displaced and stationary objects (**Figure 1C**). Vehicle-treated mice showed a statistically significant preference for the displaced object (*p* = 0.003, one-Sample t-test against 0), consistent with the acquisition, stabilization and updating of item-place memory ^37^.

In propranolol-treated animals, object exploration times also differed across sessions (Repeated-measures ANOVA; F2,23 = 15.07, *p* = 0.001), but pairwise comparisons revealed no changes between session 2 and session 3 (*p* > 0.05), with a decrease in exploration time observed only *within* session 3 compared to controls (*p* < 0.05). Between group comparisons showed significant differences between groups (Repeated-measures ANOVA; F2,28 = 15.26, *p* < 0.001), with Tukey’s post hoc tests indicating that vehicle-treated mice spent more time exploring objects than propranolol-treated mice during session 1 (*p* < 0.01), while propranolol-treated mice spent more time exploring objects during session 2 (*p* < 0.05). Moreover, propranolol-treated mice had significantly lower discrimination indices than vehicle-treated mice during session 3 (independent samples t-test; t(14) = 4.97, *p* < 0.01). The similar exploration times across session 1 and session 2, combined with the absence of a preference for the displaced item-place associations, indicate that propranolol-treated mice exhibited impairments in encoding and updating item-place associations.

### β-adrenergic receptors modulate the recruitment and reactivation of neuronal populations during item-place learning

To investigate how β-adrenergic receptors influence neuronal dynamics during spatial learning, we performed wide-field Ca^2+^-imaging of the dorsal CA1 (**Figure 2A**) using a miniature integrated microscope (miniscope) and microendoscopic GRIN lens^21^. This approach allowed us to record the activity of hundreds of neurons in freely behaving mice performing behavioral tasks (**Figure 2B**).

**Figure 2.**
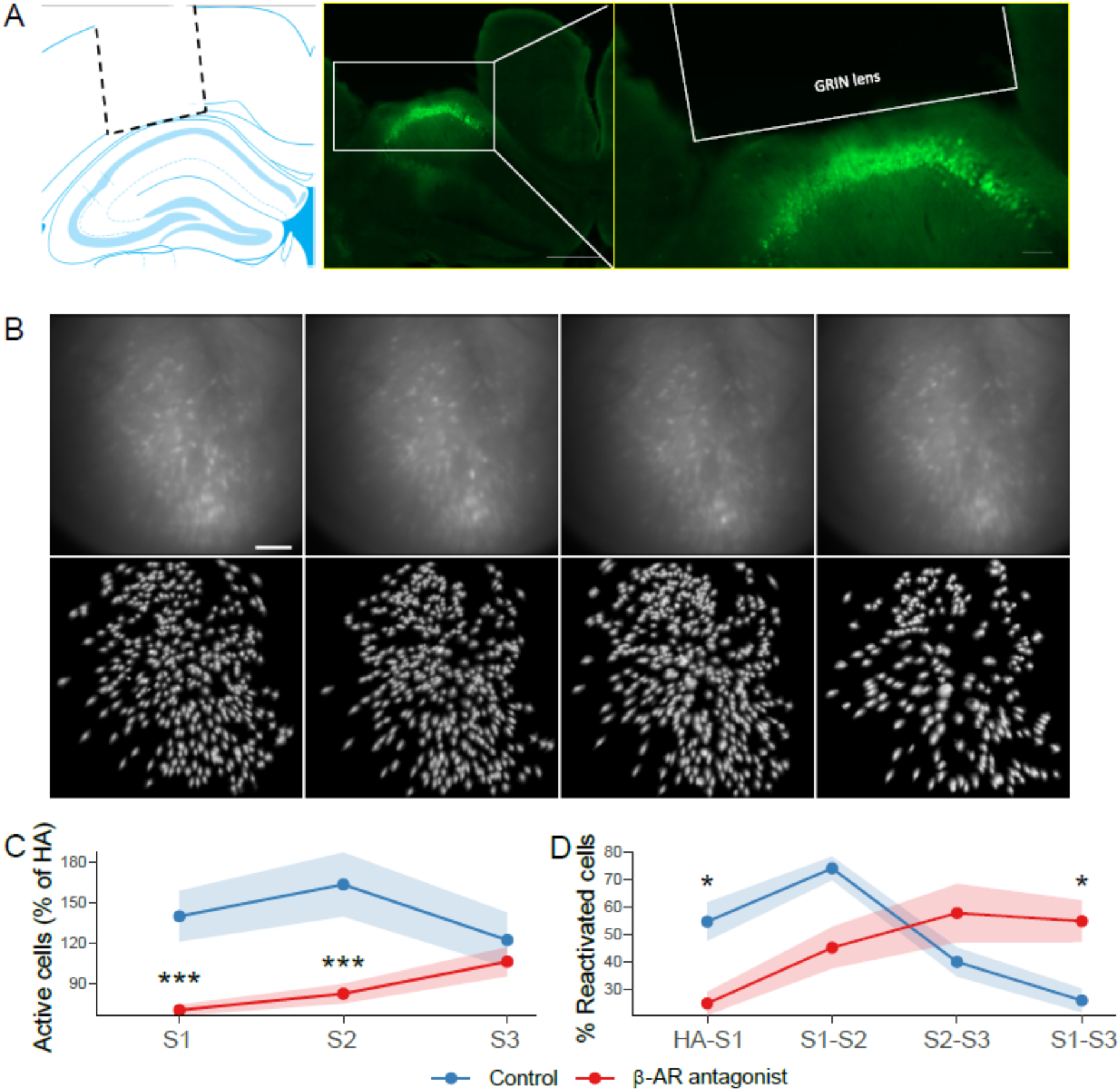
Recruitment and reactivation of neuronal populations during cumulative item-place learning. **A)** Left: Location of the GRIN lens over the dorsal CA1 (schema adapted from^38^). The dotted line shows the position of the lens over the pyramidal cell layer. Middle: white rectangle outlines pyramidal cells of the CA1 region of one mouse, that express GCaMP7f (green fluorescent label). Scale bar represents 500 μm. Right: Expansion of the same image shown in the middle panel indicating the grin lens location over the GCaMP7f-labeled cells. **B)** Top row: Raw field of view through the miniscope showing CA1 pyramidal cells detected in each session of one mouse. Left to right: habituation (HA), novel exposure (session 1, **S1**), re-exposure (session 2, **S2**) novel item configuration (session 3, **S3**). Scale bar represents 100 μmBottom row: maximum intensity projection of cell maps corresponding to the upper panels. **C)** Percentage of active cells normalized to habituation (HA) baseline values across item-place sessions S1, S2 and S3. **D)** Percentage of cells that were reactivated from one item-place session to the next, as well as between habituation (HA) and S1. Points and shaded ribbons represent mean ± SEM. Asterisks indicate statistical significance at \**p* < 0.05, \*\**p* < 0.01, \*\*\**p* < 0.001. N = 8 per group.

We first examined the recruitment of active neurons in CA1 across the task sessions (**Figure 2C**). The number of active cells per session was normalized to baseline habituation values and compared between groups. In the vehicle group, pairwise comparisons revealed no significant changes in the percentage of active cells across pairs of sessions (*p* > 0.05, Wilcoxon signed-rank test), with only a trend toward a reduced number of active cells between session 2 and session 3 (*p* = 0.08).

In contrast, propranolol-treated mice exhibited significantly less activated cells than vehicle-treated mice during sessions 1 and 2 (*p* < 0.001, Wilcoxon rank-sum test), but no difference in the percentage of activated cells was detected between controls and propranolol-treated animals during session 3 (*p* > 0.05). Within-group comparisons revealed significant changes across sessions in the propranolol group (*p* = 0.01, Friedman rank-sum test). These findings indicate that the relative percentage of neuronal participation was stable in controls across sessions, but that propranolol treatment prior to session 1, not only reduced the percentage of actively participating neurons overall, but also altered the activation dynamics across sessions.

To clarify the extent to which the population of active neurons in sessions 1 through 3 was the same neurons or different ones, we then examined the percentage of cells that were recruited in one session that were activated again in subsequent sessions (**Figure 2D**). Here, vehicle-treated mice displayed significant differences across session pairs (*p* < 0.01, Friedman rank-sum test). Specifically, they exhibited a sharp rise in neuronal reactivation *of the same neurons* between sessions 1 and 2, but showed a drop in reactivation of this population between sessions 2 and 3 (*p* < 0.01, Wilcoxon signed-rank test) and between sessions 1 and 3 (*p* < 0.001). This indicates that the reiteration of the item-place experience during session 2 recruited a similar population of neurons (that became active in session 1), but the change in spatial content in session 3, resulted in a change of ensemble properties.

The percentage of consistently reactivated cells was altered by propranolol treatment (**Figure 2D**). Here, from session to session, the percentage of re-activated cells (relative to the entire population of activated cells) steadily increased (habituation vs. session1, session 1 vs. session 2, session 2 vs. session 3, *p* < 0.05)

In summary, vehicle-treated mice re-used previously active neuronal ensembles when contexts were similar and did not preferentially recruit previous ensembles when a novel spatial content change was introduced. Propranolol-treated mice, failed to show stability in neuronal reactivation between session 1 and 2 and showed different neuronal reactivation patterns as was seen in controls during session 3.

### β-adrenergic receptor antagonism alters place cell-like properties

Despite the slower temporal dynamics of Ca^2+^ transients compared to electrophysiological recordings from neurons, wide-field Ca^2+^-imaging enables the detection of spatially responsive cells using GCAMP7 sensors^39^ - referred to here as place cell-like cells. To investigate CA1 spatial information processing in cumulative item-place learning and the role of β-adrenergic receptors in this process, we identified place cell-like cells (**Figure 3A**) and quantified the following parameters: spatial information (bits/event), to assess how well cell activity encoded spatial locations; place field size, to determine the granularity of spatial representations; spatial coherence, to evaluate the local consistency of activity patterns; and object score, which reflects the percentage of place cell-like cells encoding locations near objects^40,41^.

**Figure 3.**
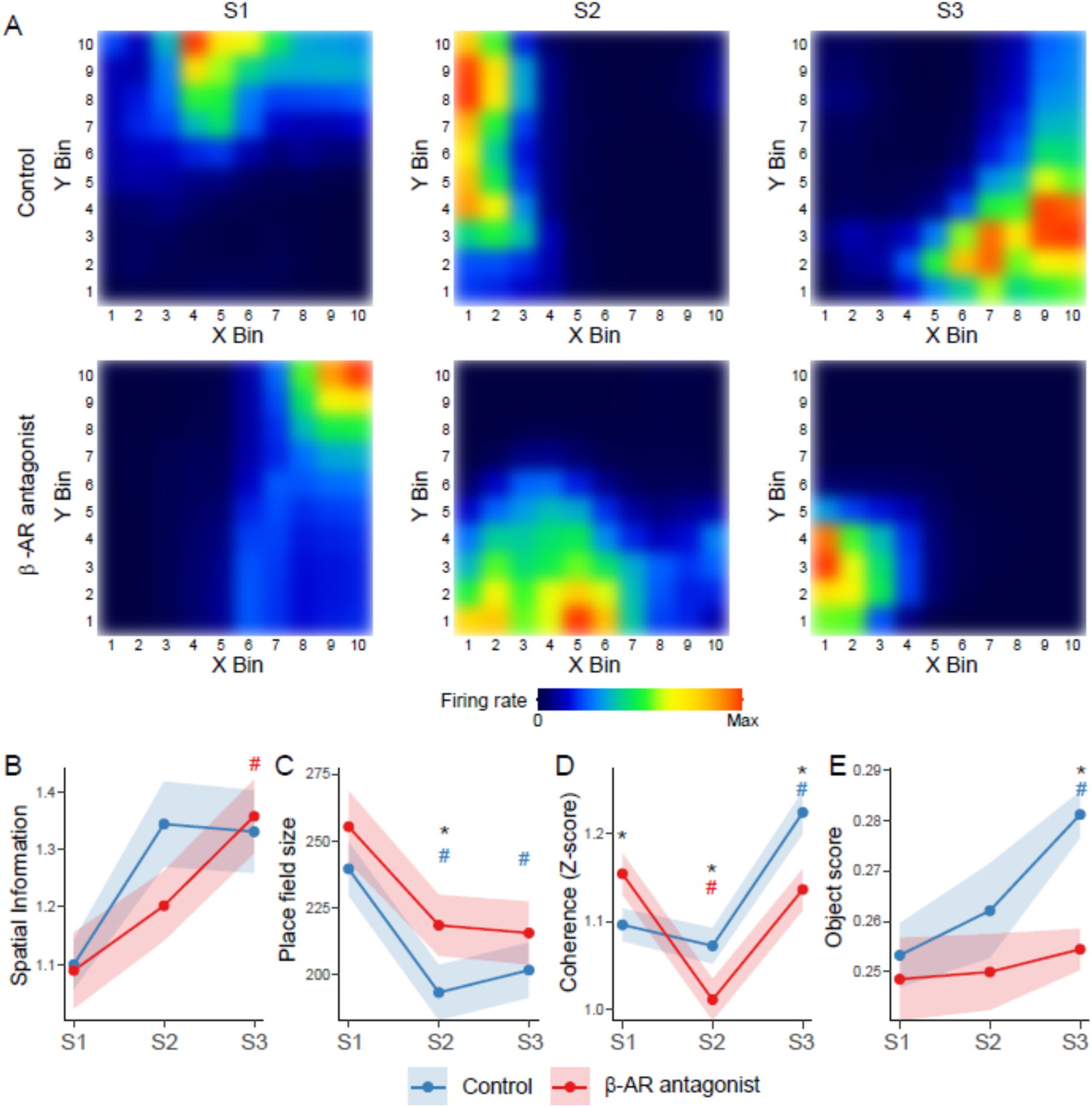
β-adrenergic antagonism alters place cell-like properties. A) Representative firing rate maps of place-cell-like cells from each group and session of the item-place learning task: **S1** (novel item-place exposure), **S2** (re-exposure), **S3** (novel item configuration). Each map depicts a different cell. **B)** Spatial Information scores (bits/event) **C)** Place field sizes (cm^2^). **D)** Z-scored coherence measure. **E)** Object scores showing average overlap of place fields with object’s vicinity. Points and shaded ribbons represent mean ± SEM. Asterisks indicate significant differencesbetween groups at \**p* < 0.05. Hashtags indicate significant within-group differences at ^#^*p* < 0.05 from S1, with blue denoting the control group and red denoting the β-AR antagonist group. N = 6 per group.

In control animals, spatial information remained stable across sessions (*p* = 0.051) (**Figure 3B**), whereas place field size showed a significant reduction (*p* < 0.001) (**Figure 3C**), with pairwise comparisons revealing a decrease from session 1 to both session 2 and session 3 (*p* < 0.05). Spatial coherence also changed significantly across sessions (*p* < 0.0001) (**Figure 3D**), remaining stable from session 1 to session 2 (*p* > 0.05), but increasing during session 3 (*p* < 0.0001). The proportion of place cell-like cells tuned to objects changed across sessions (*p* < 0.05), with a significant increase from session 1 to session 3 (*p* < 0.001) (**Figure 3E**), suggesting *de novo* integration of information about the changed item configuration. These results indicate that, in control mice, place cell-like activity in CA1 becomes more spatially refined and coherent with repeated experience and dynamically adapts subtle changes in environmental features, such as changes in spatial content. This behavior is consistent with what one would expect of electrophysiologially recorded place fields, as well as their response to spatial content change within a known environment ^42,43^.

β-adrenergic receptor antagonism resulted in different place cell-like properties across sessions. Spatial information content increased from session 1 to session 3 (*p* < 0.05) (Figure 3B), and place field size decreased from session 1 to session 2, and session 1 to session 3 (*p* < 0.05) (**Figure 3C**). The decrease in place field size was less pronounced in propranolol-treated animals than in controls during session 2 (*p* < 0.05). Spatial coherence followed a distinct trajectory (*p* < 0.0001) (**Figure 3D**), with a sharp drop from session 1 to session 2 (*p* < 0.0001), which then returned to session 1 during session 3 (*p* > 0.05), but was still lower than controls (*p* < 0.01). Unlike controls, the object scores did not change across sessions (*p* = 0.8) and were significantly lower than those of controls during session 3 (*p* < 0.0001) (**Figure 3E**).

Together, these findings suggest during item-place learning, place cell-like activity, detected via wide-field Ca^2+^ imaging in controls, becomes increasingly spatially resolved and coherent with experience, and dynamically adjusts to changes in item location. By contrast, β-adrenergic antagonism disrupts this dynamic, particularly by impairing spatial coherence and spatial tuning to the objects, potentially reflecting impaired encoding of novel spatial features /content of the environment.

### β-adrenergic antagonism dampens population bursts in CA1

Hippocampal function is characterized by population bursts - brief events during which large groups of neurons co-activate. These burst events support memory consolidation^44,45^. These events were initially described electrophysiologically, but are detectable to some extent through Ca^2+^-imaging, despite the slower temporal resolution of neuronal Ca^2+^ sensors^15,17,46^. In our recordings, CA1 neurons displayed periodic bursts of synchronized activity (**Figure 4A**). To quantify these bursts, we analyzed their number, duration, and magnitude across sessions and compared these metrics between groups.

**Figure 4.**
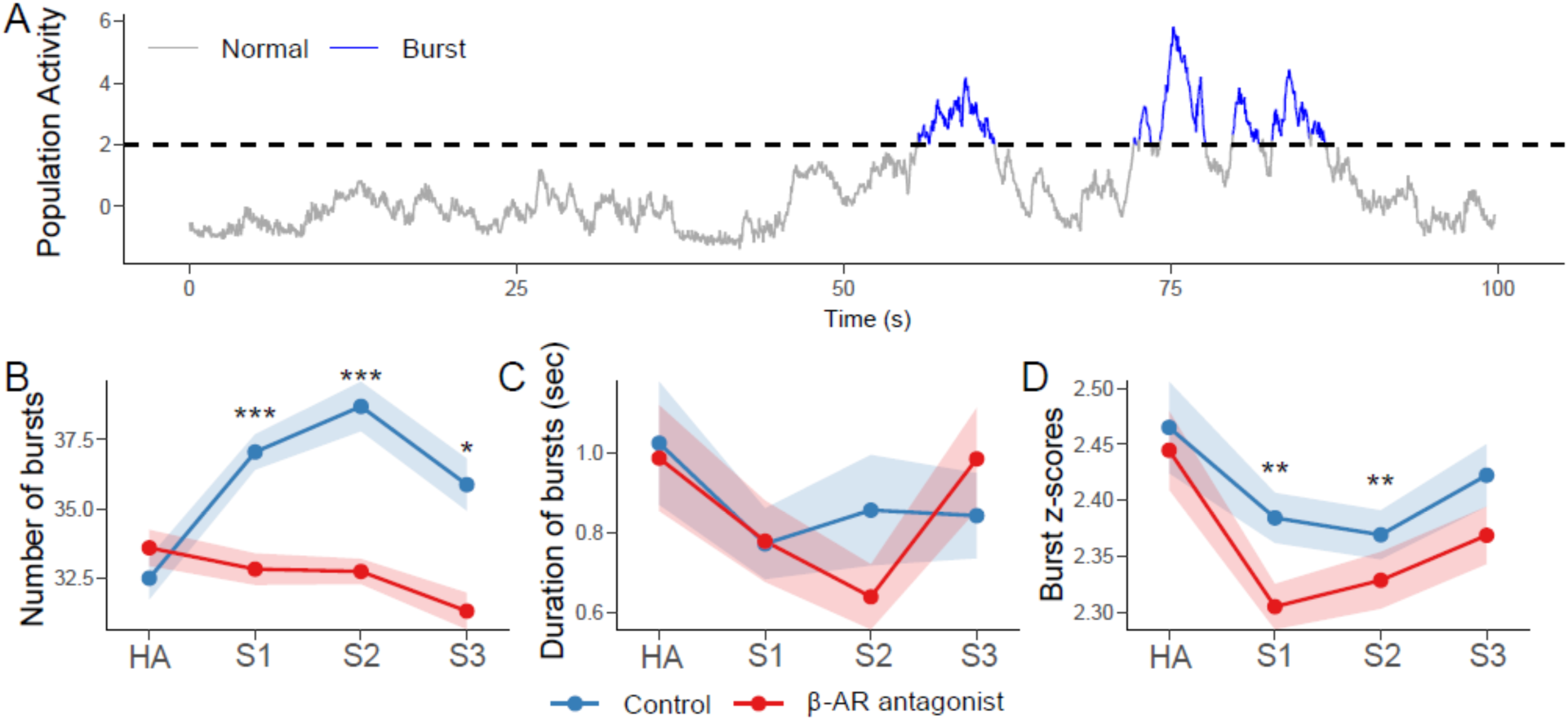
Population bursts in CA1. **A)** Z-scored mean population activity in a representative segment of CA1 recordings. The dashed line indicates the threshold (z = 2) for identifying burst events (highlighted in blue). **B)** Average number of burst events per group across sessions including habituation (HA), S1 (novel item-place exposure), S2 (re-exposure), S3 (novel item configuration). **C)** Average duration of burst events (in seconds) per group across sessions. **D)** Average z-scores of burst events per group across sessions. Points and shaded ribbons represent mean ± SEM. Asterisks indicate statistical significance at \**p* < 0.05, \*\**p* < 0.01, \*\*\**p* < 0.001. N = 8 per group.

In the vehicle group, the number of population bursts differed across sessions (*p* < 0.001, Kruskal-Wallis test) and was elevated compared to habituation during both sessions 1 and 2 (*p* < 0.001, Wilcoxon signed-rank test) (**Figure 4B**). Moreover, a slight increase was observed from session 1 to session 2 (*p* = 0.04). Both the duration and magnitude of bursts were stable across sessions (*p* > 0.05, Kruskal-Wallis test).

The antagonism of β-adrenergic receptors changed population burst activity. The number of bursts varied across sessions (*p* = 0.02, Kruskal-Wallis test) (**Figure 4B**), with pair-wise comparisons revealing a decrease during session 3 compared to habituation (*p* < 0.001, Wilcoxon signed-rank test) and lower than controls during sessions 1, 2 and 3 (*p* < 0.05, Mann-Whitney test). Burst duration remained stable and did not differ between groups (*p* > 0.05) (**Figure 4C**). However, burst magnitude, indicating the intensity of synchronized neuronal activity ^47^, changed significantly across sessions (*p* < 0.001, Kruskal-Wallis test) and was reduced during sessions 1 and 2 compared to habituation (*p* < 0.01, Wilcoxon signed-rank test), and significantly lower compared to controls (*p* < 0.01, Mann-Whitney test) (Figure 4D).

These findings indicate that β-adrenergic receptors regulate neuronal firing properties. Given the role of hippocampal bursts in supporting memory consolidation^45^, this disruption suggests that propranolol impairs CA1-mediated item-place memory encoding by dampening synchronous neuronal activity.

### β-adrenergic signaling influences network connectivity in CA1

To understand how neuronal interactions in CA1 evolve during spatial learning and how they are affected by β-adrenergic receptor blockade, we performed network analyses^48^ based on cellular calcium transients detected during the item-place learning sessions. Functional connectivity networks were constructed by calculating Spearman correlations of ΔF/F signals between all cell pairs in each session (**Figure 5A-B**). Significant positive correlations were retained to generate functional connectivity networks (**Figure 5C**), and centrality measures were calculated to capture patterns of information processing. Centrality values from subsequent sessions were normalized to the habituation (HA) baseline.

**Figure 5.**
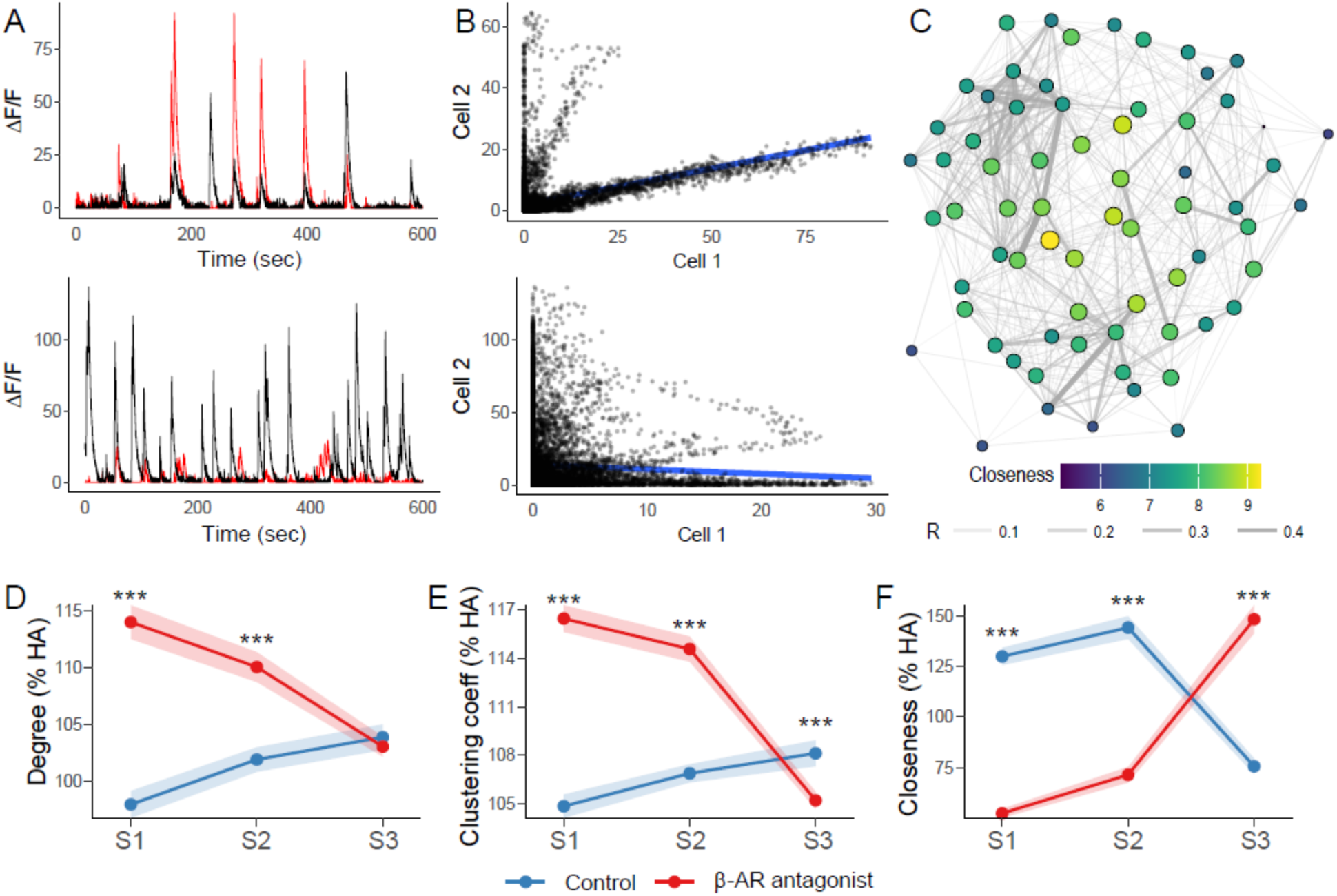
Functional network connectivity in CA1 of control animals. **A)** Example of overlaid Ca^2+^ signals (ΔF/F) from pairs of cells that were co-active (top), or not (bottom). **B)** Example of correlations between pairs of cells that were significantly correlated (top) or not (bottom). **C)** Example of a functional network of CA1 cells generated after thresholding for positive correlations (P < 0.001). Edges represent correlation coefficient values, and node size and color indicate closeness centrality. **D)** Degree centrality values expressed as a percentage of baseline values from the habituation session. **E)** Clustering coefficient values expressed as a percentage of baseline values from the habituation session **F)** Closeness centrality values expressed as a percentage of baseline values from the habituation session. S1 (novel item-place exposure), S2 (re-exposure), S3 (novel item configuration). Points and shaded ribbons represent mean ± 95% confidence intervals. Asterisks indicate statistical significance at \**p* < 0.05, \*\**p* < 0.01, \*\*\**p* < 0.001. N = 8 per group.

In control networks, ‘degree’ (**Figure 5D**), reflecting the number of synchronously active cells, changed across sessions (*p* < 0.001, Kruskal-Wallis test) and showed a progressive rise (*p* < 0.05, Wilcoxon Rank Sum test). The clustering coefficient (**Figure 5E**), reflecting the proportion of cells connected to nodes that are also interconnected, also increased (*p* < 0.001, Kruskal-Wallis test), with significant rises occurring from session 1 to session 2, and session 1 to session 3 (*p* < 0.001). The clustering coefficient remained stable, however, between sessions 2 and 3 (*p* > 0.05). Closeness centrality (**Figure 5F**), reflecting how easily a node can reach all others, was stable from session 1 to session 2 (*p* > 0.05), but dropped sharply from session 1 to session 3 (*p* < 0.001). This drop, along with the increased degree and clustering, suggests a shift towards network compartmentalization in control animals, that may be necessary to integrate novel information.

The functional connectivity of CA1 networks was profoundly changed by β-adrenergic receptor antagonism. In contrast to responses in control group, degree values decreased progressively across sessions (**Figure 5D**) (*p* < 0.05, Wilcoxon Rank Sum test). Degree was significantly higher in propranolol-treated networks compared to control networks during sessions 1 and 2 (*p* < 0.001, Mann-Whitney test), with no difference occurring during session 3 (*p* > 0.05).

Moreover, in contrast to vehicle-treated animals, clustering in propranolol-treated networks declined progressively across sessions (*p* < 0.001) (Figure 5E), and, while it was significantly higher than controls during session 1 and 2 (*p* < 0.001), it became lower during session 3 (*p* < 0.001).

Closeness values also diverged from vehicle-treated controls under propranolol treatment (Figure 5F). Closeness increased progressively across sessions (*p* < 0.001) and while it was lower than in controls during sessions 1 and 2 (*p* < 0.001) it became significantly higher during session 3 (*p* < 0.001). This progressive increase in closeness across sessions, despite decreasing degree and clustering, suggests a less specialized, more uniform network state that fails to reorganize effectively in response to novel information.

In summary, vehicle-treated networks transitioned from a sparse, efficient configuration (high closeness, low degree and clustering) to a denser, modular state (lower closeness, higher degree and clustering), in sessions 1 through 3, supporting the integration of new information, while maintaining previously encoded content. Following propranolol-treatment, networks diverged from this pattern, initially exhibiting high degree and clustering but low closeness, indicating redundancy and inefficient information flow. Over time, decreasing degree and clustering with rising closeness suggest a reliance on global connectivity without the modular organization needed to integrate spatial information effectively.

## DISCUSSION

A fundamental challenge in neuroscience is understanding how hippocampal ensembles dynamically process information during spatial learning. The dorsal CA1 region is central to these processes^14,44,49^, with noradrenergic inputs playing a key modulatory role^5,32^. Although neuronal activation patterns are known to drive memory formation, recall, and updating, the specific role of β-adrenergic receptors in orchestrating these processes remains poorly defined. Here, we employed a spatial object recognition paradigm known to engage hippocampal plasticity and to depend on β-adrenergic signaling^31^ to investigate, by means of Ca^2+^-imaging, cellular dynamics during spatial learning and memory updating. We show that cumulative item-place learning and its updating rely on progressive recruitment and reactivation of CA1 ensembles, refinement of spatially tuned neurons, consolidation-linked population bursts, and circuit connectivity changes to support the integration of new information. Moreover, we report that β-adrenergic receptors support spatial memory encoding and retrieval by modulating these processes within CA1.

In the item-place learning task, animals were exposed to the same context across three sessions: first encountering two objects (session 1), then re-encountering them in the same arrangement (session 2), and finally encountering a configuration where one object was displaced (session 3; Figure 1A). This design allows animals to encode, recall/re-iterate, and update spatial representations and thus, provides a basis for the examination of dynamic, cumulative spatial learning. Successful learning is indicated by a reduction in exploration time from session 1 to session 1, reflecting the successful acquisition of item-place memory. Memory recall and subsequent updating is indicated by the increased exploration of the displaced object during session 3. Consistent with previous work on the importance of β-adrenergic receptors in spatial learning and consolidation^33,50–52^, and particularly in associating objects with locations^31^, animals treated with the β-adrenergic receptor antagonist, propranolol failed to exhibit these behavioral indicators of learning.

Reactivation of cellular ensembles active during learning underlies memory recall^10,11^, while encoding new information and updating memories relies on integrating novel ensembles^4^. To be able to distinguish between memory acquisition, reactivation/reiteration of the memory and memory updating, we separated our sessions by 60 minutes. Animals explored in each session for 10 minutes. This duration of novel spatial exploration has been shown in the past to be an adequate time-span for the manifestation of place fields ^41^, whereas an interval of at least one hour before re-exposure to the same spatial environment results in stable place fields ^53^. Moreover, spatial learning triggers nuclear immediate early gene (IEG) expression in the CA1 region, whereby some of the neurons that were activated by the first spatial experience are recruited and/or modified by subsequent novel item-place learning ^54^.

To explore whether this kind of information storage and updating was represented by large populations of CA1 neurons and whether the detected neuronal ensembles change in an experience-dependent manner, we used the cell activity during the habituation session as a reference, and then determined how many of these cells were active in session 1. We detected a distinct population of CA1 neurons that responded to this first learning event. We observed that a large percentage of the same cells were also active in session 2, when the item-place information was repeated, suggesting re-engagement of established engram ensemble to support recall. This property of hippocampal neurons has been demonstrated using optogenetics in studies of fear memory recall ^55^ and using chemogenetic tagging of neuronal ensembles ^56^.

Reactivation of the cells that were active during sessions 1 and 2 then sharply declined in controls the spatial content (item-pace configuration) was changed during session 3 (session 2 vs. session 3 and session 1 vs. session 3), indicating a lesser engagement of previously activated ensembles when the new item-place configuration was being learned. This finding is consistent with the updating of spatial content information in CA1 neuronal ensembles^54^. In fact, re-activation of the original session 1 ensemble in its entirety might be expected to disrupt context-dependent information updating by preventing new ensemble connectivity from occurring ^57^. These interpretations are supported by our observations in animals that were treated, prior to session 1, with a β-adrenergic receptor antagonist and who failed to exhibit item-place learning, consistent with previous reports^31^.

These mice exhibited reduced ensemble reactivation, compared to controls, during the novel spatial experience of session 1 and no significant change in activity from session 1 to session 2. Interestingly, and in contrast to controls, reactivation of the neuronal ensemble increased from session 2 to 3 (between session 1 and 3). Antagonist treatment was given 30 minutes before session 1 and 170 minutes elapsed between the timepoint of injection and the beginning of session 3. This means that the antagonist is still very likely to have been effective in impairing β-adrenergic receptor function during session 3.

Taken together this pattern of neuronal activity, during β-adrenergic receptor antagonism, implies a failure of the hippocampus to distinguish between congruent and incongruent information, and suggests that these mice rely on *de novo* information encoding during session 3, rather than effectively updating representations established during sessions 1 and 2. These findings are consistent with reports that activation of β-adrenergic receptors is needed for stabilization of recently acquired spatial memory ^58^. Thus, we propose that the activation of β-adrenergic receptors is critically required for enabling the re-use of previously acquired ensembles during recall, and for recruiting new ensembles in response to novel spatial information.

Although calcium sensors do not have the temporal resolution of electrophysiological recordings, it has nonetheless been demonstrated that spatially responsive cells can be discriminated with wide-field Ca^2+^-imaging^39^. To examine the extent to which the neuronal ensemble activity we detected could align with place cell activity in the CA1 region, we examined spatial information (bits/event), place field size, spatial coherence, and object score^40,41^. In control animals we observed that place cell-like became more spatially refined and coherent from session 1 to session 2, consistent with the stabilization of place fields over time and familiarity with the spatial environment ^53,59^. Spatial coherence increased in session 3 compared to the other sessions and object scores increased. Taken together with the abovementioned finding that fewer members of the original neuronal ensemble were re-activated in session 3, this finding is consistent with rate remapping as the familiar spatial environment is updated to reflect allocentric changes in content information ^42,43^ at temporal intervals that require information retrieval and updating ^60^. To further validate these interpretations, we analyzed population bursts of neurons in the three sessions. These bursts, characterized by periodic, highly synchronous cellular activity, occur in CA1^3^ and contribute to memory consolidation and updating^16,17,61^. Bursts can be detected using wide-field Ca^2+^ -imaging despite the slower temporal resolution of neuronal Ca^2+^ sensors^15,17,46^. We observed periodic bursts of synchronized activity in the CA1 that increased in number from session 1 through 2, but decreased in session 3, whilst remaining significant from burst activity recorded during habituation. Burst firing is a characteristic of place cells ^60^. It is tempting to speculate that this change in activity during session 3 may have been driven by endogenous LTD that occurs in the CA1 region during item-place learning and information updating ^37^ and is known to modulate place fields ^62^.

During spatial object learning and updating, β-adrenergic receptor antagonism disrupted place cell-like stability and decreases spatial coherence. Findings are consistent with reports by others that locus coeruleus inputs to the hippocampus regulate place fields in a state-dependent manner^52^. β-adrenergic receptor antagonism also reduced the number of population bursts detected across sessions. These findings suggest that β-adrenergic receptors modulate burst activity in CA1, as reported for NA action in hippocampal slices^63,64^ and the antagonist-mediated disruption of burst firing may have contributing to the observed memory deficits at the behavioral level^65^. Furthermore, the altered burst dynamics likely contribute to the changes in cellular recruitment and reactivation observed across sessions.

Because brain function depends on the coordinated activity of sparse populations of neurons^9^, we also examined how β-adrenergic receptor antagonism influences CA1 circuit-level interaction. This was done by constructing functional connectivity networks^48,66,67^. In control animals, the network evolved from a sparse, efficient configuration in session 1 (low degree and clustering, high closeness) to a denser, more modular architecture during session 3 (high degree and clustering, lower closeness), indicating compartmentalization for integrating new information. In contrast, antagonist-treated networks began were dense at first, albeit inefficient at distributing information and ultimately became sparser, with information transfer efficiency peaking late in the SR task. This inverted trajectory likely hindered their ability to effectively encode novel inputs and recall previous information, further underscoring the role of β-adrenergic receptors in supporting CA1 network reconfigurations to accommodate changing demands.

In summary, our findings reveal that proper spatial learning emerges from coordinated patterns of cellular recruitment, spatial representation, and population-level dynamics within CA1. Importantly, antagonism of β-adrenergic receptors disrupts cumulative spatial learning and information updating, and alters related CA1 hippocampal ensembles at several levels. Our use of systemic propranolol treatment raises the possibility that some of these effects may stem from broader network influences, rather than direct antagonism of β-adrenergic within the CA1 region, and future research should aim to disentangle these influences and identify key modulatory regions. Collectively, these results show that to encode and update spatial memories CA1 ensembles undergo multiscale dynamic changes, from cellular recruitment to network reorganization, and that β-adrenergic receptors are essential modulators of this process.

## Author contributions

The study was designed by D M-V. Experiments were conducted by J-P W and NS. Signal analysis was conducted by NS and J-P W, and further developed by JH. Place cell and graph theory-based analysis was conducted by JH. D M-V and JH wrote the paper that was edited by all authors.

## Acknowledgements

This work was supported by a grant from the German Research Foundation (Deutsche Forschungsgemeinschaft, DFG) to Denise Manahan-Vaughan (SFB 1280/A04, project number: 316803389). We gratefully acknowledge Dr Thu-Huong Hoang and Beate Krenzek for supporting immunohistocheistry. We thank Jens Colitti-Klausnitzer his assistance in surgical interventions treatments, and Nadine Kollosch for animal care.

## Conflict of interest

The authors declare that there is no potential conflict of interest.

## Data availability

The datasets generated during and/or analysed during the current study are available from the corresponding author on reasonable request. The code and preprocessed data analyzed in this study are available at GitHub (github.com/johaubrich).

## Generative AI statement

The authors declare that no Gen AI was used in the creation of this manuscript.

## METHODS

### Animals

The study was conducted in accordance with the European Communities Council Directive for care of laboratory animals (2010/63/EU). The experiments were approved in advance by the local state authority (Landesamt für Verbraucherschutz und Ernährung, North Rhein-Westfalia). All efforts were made to minimize the number of animals used for this study, specifically by conducting power calculations to establish the minimal cohort size for meaningful statistical analyses.

Experiments were performed on eight male CBA/CaOlaHsd mice (9-11 weeks at the time of surgery; Envigo, Germany or in-house breeding). This mouse strain does not exhibit sensory deficits thoughout its lifespan, and exhibits superior spatial learning behavior and hippocampal synaptic plasticity compared to mouse strains that have chronic sensory impairments e.g. C57Bl/6 mice^68,69^. Mice were housed in in sibling groups in a temperature and humidity-controlled vivaria (Scantainer Ventilated Cabinets, Scanbur A/S, Denmark) with constant 12-hour light-dark cycle (lights on from 7 a.m. to 7 p.m.), controlled temperature (22 ± 2 °C), humidity (55 ± 5%). Food and water were available ad libitum throughout all experiments. After the surgery mice were housed individually and were allowed at least 7 day of recovery before the commencement of experiments. Experiments were performed according to the European Communities Council Directive of September 22nd, 2010 (2010/63/EU, Bezirksamt Arnsberg).

Female mice were not included they become stressed by the presence of male mice^70^ and this could alter the outcome of the behavioral experiments. For this reason, we used males only (reduction of variability of responses and keeping animal numbers low without loss of statistical power).

### Surgical procedures

Surgical procedures were adapted from methods established by others^21^. At 9-11 weeks of age, mice were anesthetized with sodium pentobarbital (60 mg/kg, intraperitoneally (i.p.)) and were head-fixed in a stereotaxic frame. After scalp hair removal, local analgesia was applied using a 10% Lidocaine spray (Xylocain® pump spray, Aspen Germany GmbH, München, Germany).

Pre- and postoperative analgesia was implemented via an s.c. injection of Meloxicam (Metacam, 0.2 mg/kg, Boehringer Ingelheim, Ingelheim am Rhein, Germany), immediately prior to an 24h after surgery.

After leveling the skull (i.e. to a horizontal plane of 0°) along the anterior-posterior (AP) axis of the skull, the same correction leveling was conducted for the medio-lateral (ML) axis, at a distance of 2 mm from the center of the line (describing the fusion of skull-bone plates) separating bregma and lambda. The depth of anesthesia was monitored during the whole surgical procedure by checking that the tail or foot pinch reflex was absent^71^.

### Ca^2+^ sensor treatment

Following the craniotomy (0.5mm in diameter), 500 nL of GCaMP7f-expressing Adeno-Associated Virus solution (PGP-AAV-syn-jGCaMP7f-WPRE, 1.8×10¹³ vg/mL, Catalogue no. 104488-AAV1, Addgene) diluted with sterile saline solution was injected through a glass micropipette (Science Products GmbH, Germany), targeting dorsal CA1 (AP: −2.1, ML: −1.4, DV: -1.3 mm with respect to bregma). Fifteen minutes after the injection, the pipette was slowly removed. The cavity was filled with bone wax (Sharpoint®,Surgical Specialties, Tijuana, Baja California, Mexico). The scalp was sutured and povidone-iodine solution (10%, Betaisodona®, Mundipharma GmbH, Frankfurt am Main, Germany) was applied over the sutured area. Ten days after this procedure, Gradient Refractive Index (GRIN) lens (part# 1050-004605, Inscopix Inc., Palo Alto, California, USA) implantation was conducted. Animals received anesthesia and analgesia as described above. The site for lens implantation was marked on the skull at AP: −2.2, ML: −2.1 mm. To limit the drilling area to the diameter of the GRIN lens, 3 locations around the craniotomy mark were additionally marked at a distance of 0.5 mm in the anterior, posterior and lateral directions. Two additional holes were drilled, one anterior to and another posterior to the left coronal suture to anchor two stainless steel screws (1 mm, Helix, Villingen-Schwenningen, Germany). Following the craniotomy (1mm in diameter), aspiration of the cortical tissue overlying the dorsal CA1 was carried out under constant supply of cold sterile physiological saline. Aspiration continued until the removal of medial-lateral striations of the corpus callosum, revealing the anterior-posterior striations. The GRIN lens (1 mm diameter, 4 mm length, product code: 1050-004623, Inscopix Inc., Palo Alto, California, USA) was implanted at a 9-degree angle over the CA1 (AP: −2.2, ML: −2.1 mm, DV: ∼ -1.4mm) under constant visual monitoring through the operation microscope (Model OPMI 1-FC, Carl Zeiss, Oberkochen, Germany). Once the GRIN lens was implanted, the physiological saline, solution located in the gaps between the skull edge and the GRIN lens, was removed and replaced by cyanoacrylate adhesive (UHU® Sekundenkleber, UHU GmbH & Co KG, Bühl, Germany). Acrylic dental cement (Paladur, Heraeus Kulzer GmbH, Hanau, Germany) was further applied onto the dried cyanoacrylate adhesive, and around the screws affixed to skull. Fourteen days after lens implantation mice were anesthetized with sodium pentobarbital (60 mg/kg, i.p.) and their GCaMP signal was confirmed using a miniscope (Inscopix Inc., Palo Alto, California, USA) set at the middle of its focal range and attached to a baseplate (product code: 1050-004638 Inscopix Inc., Palo Alto, California, USA). The miniscope was connected to an nVista 2.0 imaging system (Inscopix Inc., Palo Alto, California, USA). After acquiring an in-focus view of dorsal CA1 region, the baseplate was glued to the skull using dental cement (Paladur® Heraeus Kulzer GmbH, Hanau, Germany). Parameters associated with miniscope imaging such as focus level, excitation light intensity, and gain were also finalized. At 3-4 days prior to the recording sessions, mice were handled by the experimenter and habituated for a minimum of 20 min, twice a day to the procedure of attaching the miniscope. This was carried out in the same experiment room where the imaging chamber was located, in order to habituate the mice to the experiment environment. The same experimenter handled all mice during all handling and experiment days.

### Item-place Learning Task

We used a cumulative item-place task to examine the ability of the mice to distinguish between an object that is located at a familiar location or an object that is spatially novel^31^. The task took place in a 40 x 40 x 50 cm arena. Item-place experiments were performed in three 10 min sessions separated by 60 min intervals (Figure 1A). On day 1, Ca^2+^ signals were recorded at 20 FPS while mice explored the arena without in the absence of any physical items (habituation). On day 2, during session 1 (novel exploration), two novel objects were presented for exploration. The mice were then re-exposed to the objects in the same locations in the next session (session 2), and in the third session (session 3) one object was shifted in a parallel fashion to the other side of the arena to test for item-place recall and information updating. Objects as well as their relative positions were randomly assigned for each animal. The imaging chamber was cleaned thoroughly between task trials to ensure the absence of olfactory cues. After every trial and before the first presentation, the objects were cleaned with ethanol, rinsed with water and dried. The objects were distinctly different from one another and heavy enough so that they could not be moved by the mice. Several copies of each object were available.

Behavioral data for the item-place experiments were recorded from a camera (acA-1300, Basler AG, Ahrensburg, Germany) using EthoVision XT software (Noldus Information Technology, Wageningen, The Netherlands). Exploration of the objects was then analyzed post-hoc using the within-object area scoring system which was defined as sniffing of the object (with nose contact or head directed to the object) within ∼2 cm radius of the object^72^. Standing, sitting or leaning on the object was not scored as object exploration.

### Pharmacological Treatment

The competitive β-adrenergic receptor antagonist, propranolol hydrochloride (Tocris Bioscience, Abingdon OX14 3NB, United Kingdom), was dissolved in 0.9 % NaCl solution and administered intraperitoneally (i.p.) at a dose of 20 mg/kg and at a volume of 0.01 ml/g, and applied 30 min before novel exposure session 1, as described previously^31^. Control animals were injected with vehicle solution (0.9 % NaCl, 0.1 ml/g).

### Signal analysis

Extraction of Ca^2+^ signals from individual neurons was carried out using Inscopix Data Processing Software (IDPS) (Inscopix, Palo Alto, CA, USA). Preprocessing stages involved (i) selecting the region of interest (cropping), (ii) spatial down sampling by a factor of two, (iii) spatial filtering and (iv) motion correction. Putative cells were identified using ‘constrained non-negative matrix factorization for microendoscope data’ (CNMFe) toolbox^73^. Signal from individual cell in each cell-set was then carefully checked through visual curation to make accept/reject decision based on location of signal relative to the cell contour and corresponding appearance of peak in the signal time trace. Only data from cells labelled as ‘accepted’ were used to create spatial footprints needed for alignment and longitudinal cell registration in CellReg toolbox as described by Sheintuch and colleagues^74^. The habituation session was treated as the reference session, as it provides a scaffolding network over which further experiences during the item-place task were built. To compare active cells between the sessions, cell count in each session was normalized to habituation session count and expressed as percentage value. Reactivation of cells in a pair of successive sessions was also computed. For this, the percentage of cells in a given session that were reactivated in the subsequent session was calculated for each pair of sessions as shown in Fig. 2D Differences between groups were determined by Mann-Whitney U tests. Within-group comparisons were calculated using Friedman rank sum test followed by Wilcoxon signed-rank test.

Bursts were identified by first z-scoring the calcium traces of all individual neurons recorded in a session and then computing the population mean activity at each time point. This population signal was subsequently z-scored, and periods exceeding a threshold of z = 2 were considered candidate bursts. To define a burst event, we marked the onset when the activity crossed above the threshold and continued to hold above it until it fell below the threshold again. Bursts shorter than 0.05 seconds were discarded to exclude transient fluctuations. Differences between groups were determined by Mann-Whitney U tests.

For functional network analysis, data containing changes in cellular fluorescence intensity were used. For each animal and session, pairwise Spearman correlations with Bonferroni correction between all cells were computed and only correlations with p values < 0.001 were considered as ‘edges’ in the resulting network. Different measurements of centrality were calculated to elucidate the connectivity patterns of the networks^57,75,76^. The degree of each node was calculated as its number of edges and was normalized by the overall network size. The clustering coefficient of each node estimated the likelihood of its neighbors being interconnected and incorporated the edge weights. Closeness centrality of each node was defined as the inverse of its average distance to all other nodes in the network, also taking into account the edge weights (r values). Differences between groups were determined by Mann-Whitney U tests. Within-group comparisons were calculated using Kruskal-Wallis test followed by Wilcoxon signed-rank test.

### Identification of place cell-like activity

Raw x-y positions extracted from the tracking software were converted from pixels to centimeters and down-sampled to a temporal resolution of 0.5 seconds. Denoised, deconvolved calcium transients were treated as spike events. The arena was divided into 10 x 10 bins (∼ 4 cm x 4 cm each) and each spike event was assigned based on its coordinates. Bins without events were filled with zeros, and the proportion of spatial bins visited (occupancy) was calculated.

Raw firing-rate maps were generated for every recorded neuron. Rate maps were denoised by convolving both the spike histogram and the occupancy histogram with an isotropic Gaussian kernel of width *σ* = 1. Spatial selectivity was measured with the Skaggs information metric as previously described^67^. Statistical significance was assessed by a circular-shuffling procedure: for each neuron the entire event train was rotated in time by a random offset while the behavioral record remained fixed, and the information score was recomputed 500 times. The empirical p-value or a neuron was the proportion of shuffled scores that equaled or exceeded the observed score; neurons with P < 0.05 were retained for further analysis. To ensure adequate sampling, neurons displaying fewer than 20 events during the recording were rejected. A cell was finally classified as a place cell-like cell only if it satisfied both criteria: significant spatial information (P < 0.05) and a minimum of 20 calcium events.

The spatial extent of individual place fields was identified on each smoothed rate map by thresholding at 20 % of the peak firing rate, labelling only contiguous above-threshold regions, as described by Harvey and colleagues^77^. For every accepted field the centroid coordinates and field size were recorded, and multi-field were allowed.

To calculate spatial coherence^40,41^, after Gaussian smoothing, the unsmoothed rate matrix and its box-car–smoothed counterpart were vectorized and their Pearson correlation coefficient was obtained over all spatial bins that contained at least one sample. A Fisher’s transform was applied, yielding a coherence score that increases with the spatial contiguity of firing.

To detect place-cell-like cells tuned to bins containing objects, a neighborhood extending two bins beyond each object’s centroid bin in every direction was defined, and a mask was created. To quantify how strongly a field overlapped an object region an “object score” was calculated. The thresholded smoothed rate maps were intersected with the padded object mask, and a ratio of overlap was calculated, ranging from 0 (no overlap) to 1 (the field entirely was contained within an object zone).

### Statistical analysis

Behavioral data were analyzed using either repeated-measures ANOVA followed by Tukey’s post hoc tests or independent samples t-tests. Object exploration indexes were also compared against 0 using a one-sample t-test.

The Shapiro–Wilk test indicated that data related to cellular recruitment and reactivation, population bursts, place cell metrics, and network properties were often non-normally distributed. Therefore, non-parametric statistical tests were used for both within-group and between-group comparisons. Kruskal–Wallis tests were used to detect overall differences across sessions. When significant, these were followed by Wilcoxon rank-sum tests for within-group comparisons across sessions. Mann–Whitney U tests were used for pairwise between-group comparisons. The p-values were adjusted for multiple comparisons using the Holm method.

All tests were two-tailed, and significance was set at p < 0.05.

These analyses were conducted using R (version 4.2.2) using the R packages igraph (https://igraph.org/r), rstatix (https://CRAN.R-project.org/package=rstatix) and custom code written in R available at https://github.com/johaubrich.

## Notes

### Competing Interest Statement

The authors have declared no competing interest.

