## Supplementary Figure 1 for "β-adrenergic receptors modulate CA1 population coding during cumulative spatial memory formation and updating"

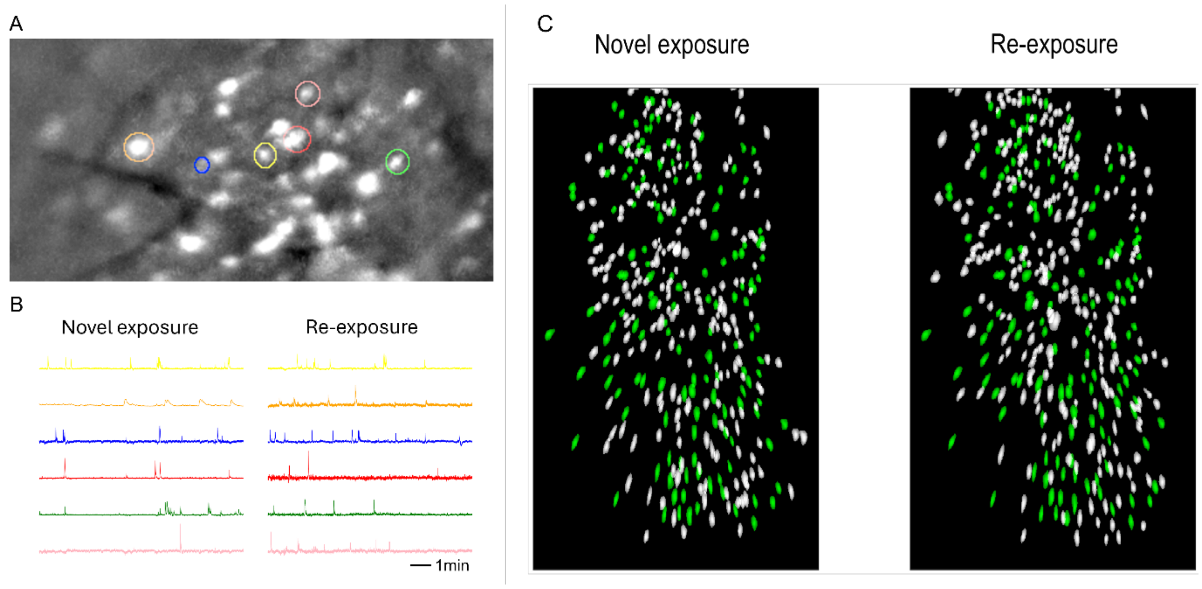


**Supplementary Figure S1**:

(A) Contours show fluorescence activity from example cells that were detected in two subsequent sessions - novel exposure and re-exposure.

(B) Raster plots of detected neuronal activity of cells highlighted in A.

(C) Spatial footprints of cells detected in novel item-place exposure (session 1, left) and re-exposure (session 2, right) with cells reactivated in both sessions highlighted in green.
